# Improving Martini 3 for disordered and multidomain proteins

**DOI:** 10.1101/2021.10.01.462803

**Authors:** F. Emil Thomasen, Francesco Pesce, Mette Ahrensback Roesgaard, Giulio Tesei, Kresten Lindorff-Larsen

## Abstract

Coarse-grained molecular dynamics simulations are a useful tool to determine conformational ensembles of proteins. Here, we show that the coarse-grained force field Martini 3 underestimates the global dimensions of intrinsically disordered proteins (IDPs) and multidomain proteins when compared with small angle X-ray scattering (SAXS) data, and that increasing the strength of protein-water interactions favours more expanded conformations. We find that increasing the strength of interactions between protein and water by ca. 10% results in improved agreement with the SAXS data for IDPs and multi-domain proteins. We also show that this correction results in a more accurate description of self-association of IDPs and folded proteins and better agreement with paramagnetic relaxation enhancement data for most IDPs. While simulations with this revised force field still show deviations to experiments for some systems our results suggest that it is overall a substantial improvement for coarse-grained simulations of soluble proteins.

## Introduction

Intrinsically disordered proteins (IDPs) are proteins that do not fold into a single well-defined structure, but rather sample a range of conformations (***Wright and Dyson, 1999***). Similarly, multidomain proteins consisting of folded domains connected by flexible linkers or intrinsically disordered regions (IDRs) are conformationally dynamic, as the folded domains can reorient with respect to each other (***Delaforge et al., 2016***). Molecular dynamics (MD) simulations are a useful tool for structural characterization of IDPs and multidomain proteins. Using integrative methods, MD simulations can be used to determine conformational ensembles of IDPs and multidomain proteins in accordance with experimental data (***Thomasen and Lindorff-Larsen, 2022***). Successful application of MD simulations relies on accurate force fields and adequate sampling of protein conformations (***Bottaro and Lindorff-Larsen, 2018***).

Coarse-grained MD simulations, where groups of atoms are represented by single beads, allow for efficient sampling of IDP and multidomain protein conformations (***Ingólfsson et al., 2014***). One of the most widely used coarse-grained force fields for biomolecular systems is Martini (***Marrink et al., 2007; Monticelli et al., 2008***). Martini maps two to four non-hydrogen atoms to one bead and is mainly parameterized against thermodynamic partitioning data. While Martini has been used successfully to study a wide range of biomolecular systems, earlier versions of the force field have been found to underestimate the global dimensions of flexible multidomain proteins (***Larsen et al., 2020; Martin et al., 2021; Jussupow et al., 2020***) and overestimate protein-protein interactions (***Stark et al., 2013; Berg and Peter, 2019; Alessandri et al., 2019; Benayad et al., 2021; Majumder and Straub, 2021; Lamprakis et al., 2021***). In order to favor more expanded conformations of multidomain proteins, we have previously used an approach based on increasing the strength of non-bonded interactions between protein and water beads (***Larsen et al., 2020; Martin et al., 2021***), improving the agreement with SAXS experiments. Similarly, others have decreased the strength of non-bonded interactions between protein beads to improve the accuracy of IDP phase partitioning (***Benayad et al., 2021***) and protein-protein interactions (***Stark et al., 2013***).

A new version of the Martini force field, Martini 3, was recently released, featuring a rebalancing of non-bonded interaction terms and addition of new bead types (***Souza et al., 2021***). Martini 3 shows improved accuracy for a wide range of systems in biology and materials science and a high level of transferability. Improved areas include molecular packing, transmembrane helix interactions, protein aggregation, and DNA base-pairing (***Souza et al., 2021; Lamprakis et al., 2021***). Here, we have tested the ability of Martini 3 to reproduce the global dimensions of IDPs and multidomain proteins. We find that simulations with Martini 3 on average underestimate the radius of gyration (*R_g_*) by ≈ 30%, and suggest a rescaling factor for increased protein-water interactions that improves agreement with small angle X-ray scattering (SAXS) data and alleviates problems with overestimated protein-protein interactions.

## Results and Discussion

We chose a set of twelve IDPs and three multidomain proteins to cover a range of different systems with available SAXS data (***Riback et al., 2017; Cordeiro et al., 2019; Mylonas et al., 2008; Riback et al., 2017; Ahmed et al., 2021; Martin et al., 2020; Johnson et al., 2017; Gomes et al., 2020; Kjaergaard et al., 2010; Jephthah et al., 2019; Fagerberg et al., 2020; Sonntag et al., 2017; Martin et al., 2021***) and ran MD simulations of each protein using the Martini 3 force field. For all proteins, we found that the ensemble generated with Martini 3 was too compact when comparing the average *R_g_* from the simulation with the *R_g_* calculated from SAXS data using Guinier analysis (Fig. 1a-b, e). A direct comparison with the experimental SAXS data also revealed deviations beyond the level expected by experimental errors (Fig. 1 c-d).

**Figure 1.**
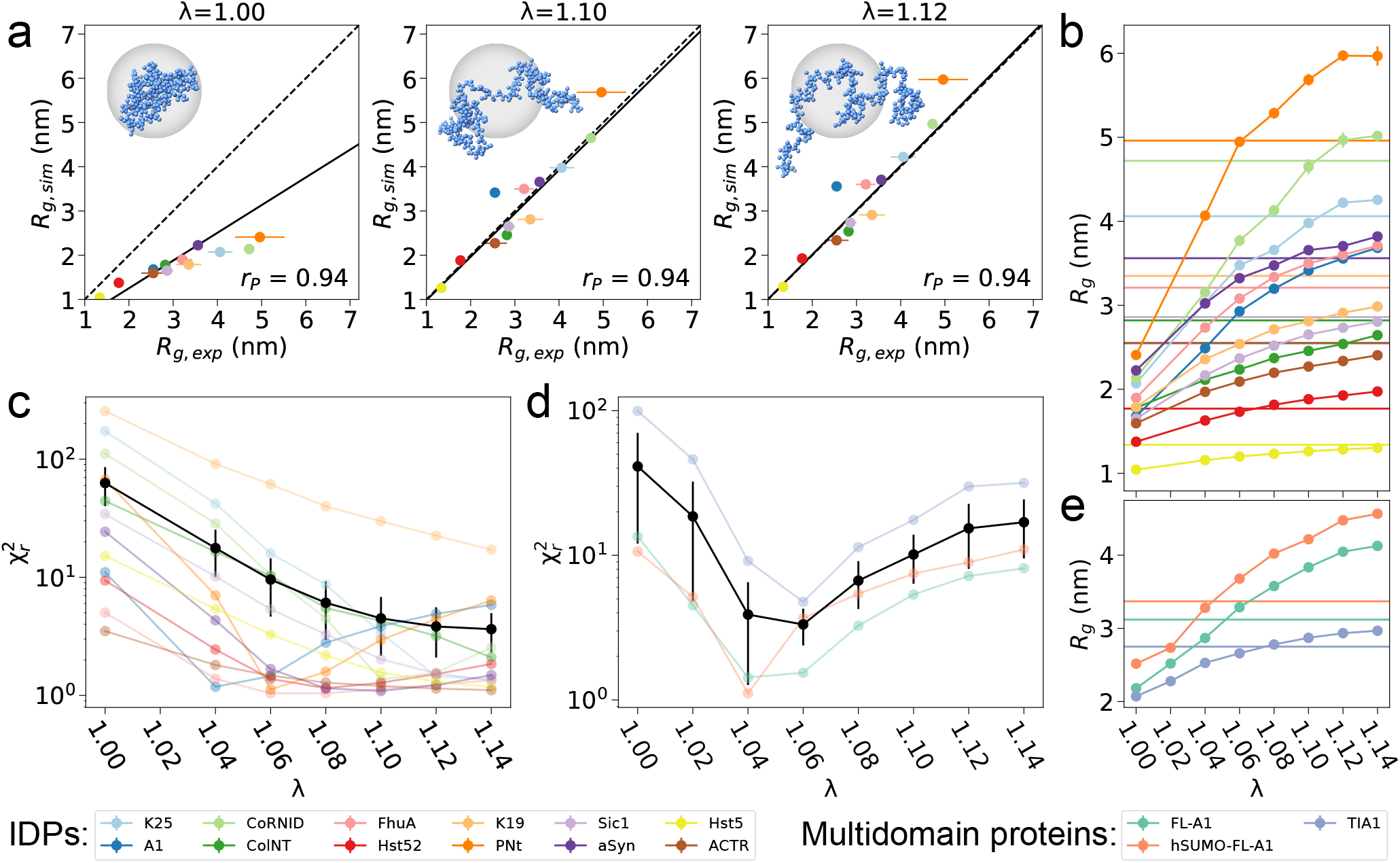
Increased protein-water interactions improve the agreement with SAXS data for IDPs and multidomain proteins. **a.** Average *R_g_* from MD simulations with three different rescaling factors for the protein-water interactions (*λ*) plotted against experimental *R_g_* from Guinier analysis of SAXS data fora set of twelve IDPs. Error bars for the experimental values were determined in the Guinier fit, and those for the simulations (here and elsewhere) were determined by block error analysis (***Flyvbjerg and Petersen, 1989***). A linear fit with intercept 0 weighted by experimental errors is shown as a solid line. The Pearson correlation coefficient (*r_P_*) is shown. The insert shows structures of Tau K25 (***Mylonas et al., 2008***) with the average *R_g_* found for each *λ*. See Fig. S1 for similar plots for other values of *λ*. **b.** Average *R_g_* from MD simulations over a range of *λ*-values for a set of twelve IDPs. Experimental values from Guinier analysis of SAXS data are shown as horizontal lines. **c–d.** Reduced 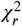 between SAXS profiles calculated from MD simulations and experimental SAXS profiles for a range of *λ*-values for a set of twelve IDPs (c) and three multidomain proteins (d). Average 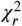 is shown in black with standard error of the mean as error bars (note the log scale). **e.** Average *R_g_* from MD simulations over a range of *λ*-values for three multidomain proteins. Experimental values from Guinier analysis of SAXS data are shown as horizontal lines. Data and scripts are available via http://github.com/KULL-Centre/papers/tree/main/2021/Martini-Thomasen-et-al

For atomistic force fields, it has previously been shown that increasing the protein-water interactions will favour expanded conformations of IDPs, resulting in more accurate global dimensions (***Best et al., 2014***). Inspired by this approach, we increased the strength of protein-water interactions by rescaling the *ϵ* parameter in the non-bonded Lennard-Jones potentials between all protein and water beads by a rescaling factor, *λ*. For all proteins, increased protein-water interactions (*λ*>1) resulted in an increased *R_g_* and improved agreement with SAXS data as measured by the reduced 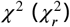. To determine an optimal value of *λ* for IDPs, we scanned six *λ*-values from 1.04 to 1.14 for each IDP. Based on the 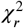 to SAXS data and agreement between *R_g_* calculated from ensemble coordinates and *R_g_* calculated from experimental SAXS profiles we chose *λ*=1.10 as a balance between improving agreement with experiments and keeping the force field as close as possible to the original (Fig. 1a-c). We performed the same analysis for the three multidomain proteins with flexible linkers, including also *λ*=1.02. These had optimal values of *λ* around 1.06 (Fig. 1 d-e), suggesting that the optimal value may be different for folded domains and IDPs.

We examined whether the too compact IDP structures in Martini 3 could be amended by simpler changes to the simulation setup instead of rescaling the protein-water interactions. To test whether including long-range electrostatics would favor more expanded conformations, we ran simulations of Histatin-5 and α-synuclein with Particle Mesh Ewald (PME) electrostatics (Fig. S2). The radii of gyration were, however very similar to those obtained using the standard reaction-field method, with some small differences for the longer protein (α-synuclein) when it was more expanded. To examine whether a lack of transient secondary structure in the simulations compared to experiments caused the compaction, we ran simulations of ACTR restraining it to form a helical structure in two regions that transiently sample helices in solution (***Kjaergaard et al., 2010***). This did not solve the problem either (Fig. S2). Finally, to examine whether the observed IDP compaction was a result of a lack of bulk solvent in our simulations, we ran simulations of α-synuclein in a very large box (*d* = 34.1 nm), but the results were essentially the same as in the smaller (*d* = 24.1 nm) box (Fig. S2). Given that the compaction of the IDPs was not substantially affected by these changes, we continued with our approach of increasing protein-water interactions.

To further investigate the effect of rescaling protein-water interactions, we performed a number of tests comparing the original force field (*λ*=1) to the force field with increased protein-water interactions (*λ*=1.10 and 1.12). First, we tested the effect on the intrachain interactions in IDPs by comparing paramagnetic relaxation enhancement (PRE) data calculated from simulations of α-synuclein, the FUS low-complexity domain (LCD), the LCD in hnRNPA2 (A2), full-length tau (htau40), and osteopontin (OPN) to PRE experiments (***Dedmon et al., 2005; Monahan et al., 2017; Ryan et al., 2018; Mukrasch et al., 2009; Platzer et al., 2011***). We found that increasing the strength of proteinwater interactions improved the agreement with PRE data as quantified by the 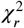 for all proteins except A2 LCD (Fig. 2a).

**Figure 2.**
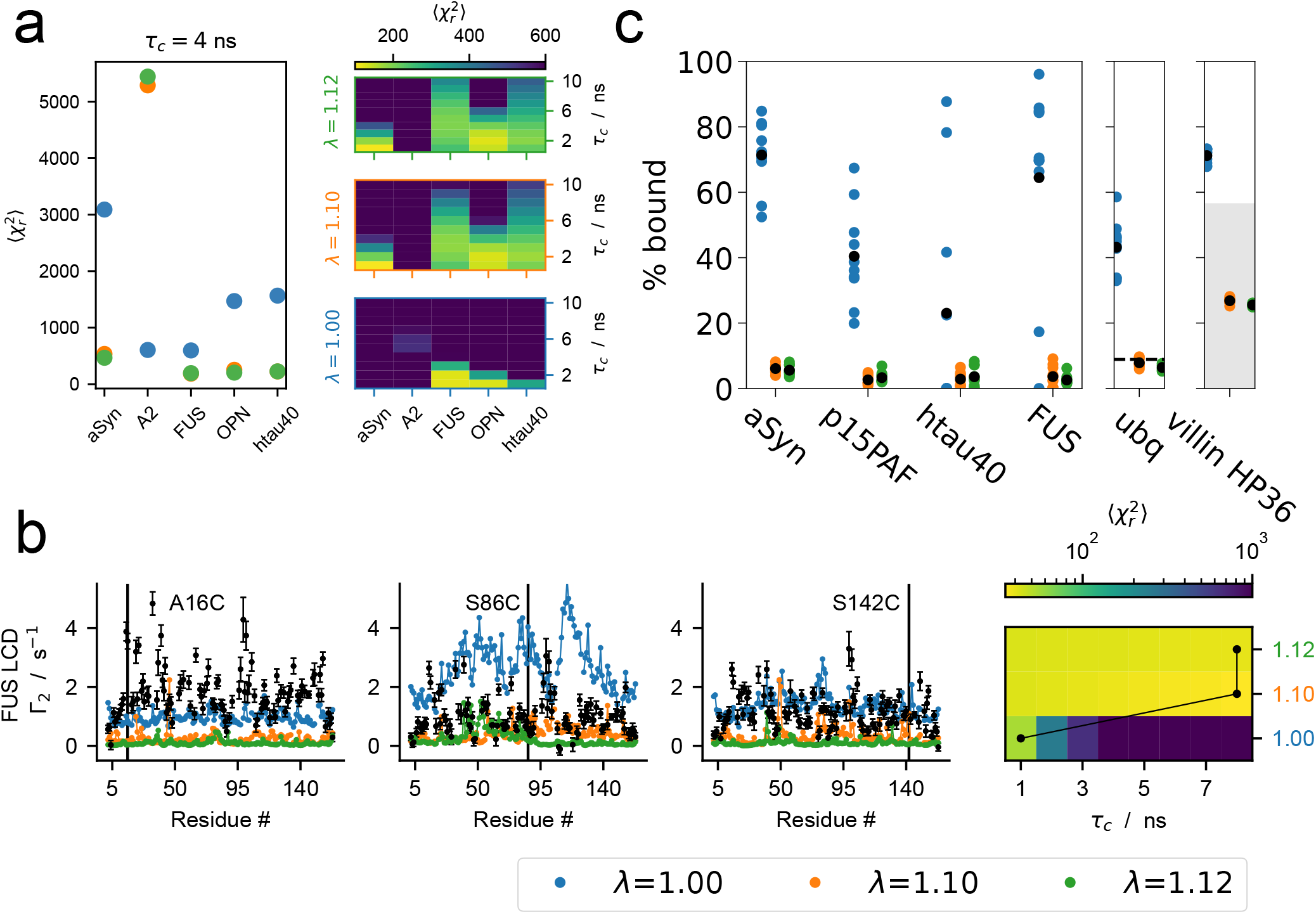
Effect of increased protein-water interactions on intrachain contacts and protein-protein interactions. **a.** Agreement between intrachain PREs calculated from MD simulations with different protein-water interaction rescaling factors *λ* and experimental PREs for the five IDPs α-synuclein, A2 LCD, FUS LCD, OPN, and htau40 measured by 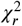. Left panel shows results with *τ_c_* =4 ns. Right panel shows that the results are consistent across a range of *τ_c_*-values (see also Figs. S3 and S4). **b.** Interchain PREs calculated from MD simulations with different *λ*-values of two copies of FUS LCD and comparison with experimental PREs (black). PREs are shown for three spin-label sites marked with black lines. Rotational correlation time *τ_c_* was selected individually for each *λ* to minimize 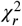. For results at *τ_c_* = 6 ns, see Fig. S5 **c.** Fraction bound calculated from MD simulations of two copies of the IDPs α-synuclein, p15_*PAF*_, htau40, and FUS LCD and the folded proteins ubiquitin and villin HP36 with different protein-water interaction rescaling factors *λ*. The results from ten replica simulations are shown as colored points with the average value shown in black. The fraction bound in agreement with *K_d_* =4.9mM for ubiquitin self-association is shown as a dashed line (***Liu et al., 2012***). The fraction bound in agreementwith a *K_d_* >1.5mM for villin HP36 self-association is shown as a shaded gray area (***Brewer et al., 2005***).

Next, we tested the effect of rescaling protein-water interactions on interchain IDP-IDP interactions. We simulated two copies of the FUS LCD at conditions matching interchain PRE experiments (***Monahan et al., 2017***) and calculated interchain PREs from the simulations for comparison. Again, increasing protein-water interactions improved the agreement with PRE data (Fig. 2b). These results suggest that increasing the strength of protein-water interactions results in more accurate intra- and interchain interactions for IDPs.

As a negative test of IDP-IDP interactions, we ran simulations with two copies of IDPs that were not expected to interact substantially. We chose α-synuclein, htau40, and p15_*paf*_, which should not interact under the given simulation conditions based on experimental evidence (***Dedmon et al., 2005; Mukrasch et al., 2009; Platzer et al., 2011***). Our results show that the original force field substantially overestimated IDP-IDP interactions, predicting a high population of the bound state. For all proteins, increasing protein-water interactions by *λ*=1.10 and 1.12 reduced the population of the bound state to below 10%, improving the agreement with experiment (Fig. 2c). We note that it was not in all cases possible to converge the populations of the bound and unbound states in simulations with the unmodified force field, as the complexes stayed bound for very long (examples in Figs. S6 and S7). However, our simulations were started from the unbound state, so we expect that a lack of convergence would result in underestimation of the bound state population at *λ*=1. For *λ*=1.10 and 1.12, several binding and unbinding events were sampled, and the distribution of the fraction bound over independent simulations was narrower (Fig. 2c).

For comparison, we also calculated the population of the bound state in our simulations of the FUS LCD, which should associate to a measurable extent based on PRE experiments (***Monahan et al., 2017***). However, FUS LCD had a population of the bound state similar to the IDPs that should not self-associate (Fig. 2c). Since the agreement with interchain PREs was improved for FUS LCD after increasing protein-water interactions, it may be that the bound state population of FUS LCD is accurate, while the bound state of the non-interacting IDPs is still slightly overestimated after increasing protein-water interactions. Nevertheless, the affinities of IDP self-association are much improved.

Finally, we investigated the effect of rescaling protein-water interactions on interactions between folded proteins. Inspired by previous simulations (***Best et al., 2014; Berg and Peter, 2019***) and nuclear magnetic resonance (NMR) experiments (***Brewer et al., 2005; Liu et al., 2012***), we simulated two copies of the villin headpiece HP36 (villin HP36) and two copies of ubiquitin, and calculated the populations of the bound state (Fig. 2c). Simulations with Martini 3 appeared to overestimate ubiquitin homodimerization when compared with the dissociation constant *K_d_* = 4.9±O.3 mM determined by NMR chemical shift perturbations (***Liu et al., 2012***), but rescaling the strength of protein-water interactions by *λ*=1.10 improved the agreement with the experimental affinity. We note that the interactions observed in the simulations were not specific to the homodimerization site determined by NMR (***Liu et al., 2012***) (Fig. S8). Although salt-dependent aggregation behavior was shown to be qualitatively improved for villin HP36 in the Martini 3 publication (***Souza et al., 2021***), homodimerization of villin HP36 also appeared to be overestimated with the unmodified force field. Based on diffusion coefficients determined by NMR, villin HP36 homodimerization should have a *K_d_*>1.5 mM (***Brewer et al., 2005***), but the population of the bound state was higher than expected for this affinity. After rescaling protein-water interactions by *λ*=1.10 and 1.12, the populations of the bound state were in agreement with *K_d_*>1.5 mM. Thus, increased protein-water interactions also seem to improve the affinities of protein-protein interactions for folded proteins, although the lack of specificity in these interactions may skew the results, as observed for ubiquitin.

## Conclusions

Our results show that simulations with the Martini 3 force field result in underestimated global dimensions of IDPs and multidomain proteins, and that rescaling the Lennard-Jones potentials for protein-water interactions by a factor *λ*=1.10 improves agreement with SAXS experiments. Additionally, this improves the agreement with PRE data for all but one of the tested IDPs, suggesting improved accuracy of intra- and interchain interactions. Our results also show that Martini 3 greatly overestimates IDP homodimerization, indicating that IDP-IDP interactions are too strong, but increasing protein-water interactions leads to a more accurate balance. The same is true for homodimerization of the folded proteins ubiquitin and villin HP36. For multidomain proteins containing flexible linkers or IDRs, a rescaling factor of *λ*=1.06 seems to be sufficient to result in good agreement with the SAXS data, although this is based on a smaller set of proteins. We note that the agreement with experiments at *λ*=1.10, determined as the optimal value for IDPs, is also better than at *λ*=1 for multidomain proteins.

For systems with no additional information available, we suggest setting *λ*=1.10. If one wishes to modify the original force field as little as possible, *λ* can bet set to 1.06 for multidomain proteins, although *λ*=1.10 shows improvement over the original Martini 3 force field for all systems tested here and may be a good overall compromise across different systems. As an alternative approach, a *λ*-value can be chosen specifically for the system of interest if the level of compaction has been probed experimentally. However, this does not necessarily entail optimizing the *λ*-value for every condition of interest. For example, we have previously selected a single *λ*-value for simulations of full-length hnRNPA1 (A1) (with a beta version of Martini 3) based on SAXS data at one salt concentration, and studied the effect of salt on the level of compaction by keeping the *λ*-value fixed but varying the salt concentration (***Martin et al., 2021***). A similar approach may be useful to transfer *λ*-values between proteins with related sequence properties, for example in mutagenesis studies.

We note that our approach of rescaling protein-water interactions does not perfectly capture sequence-dependent differences in IDP compaction. This is evidenced by the observation that the optimal value of *λ* correlates with the relative expansion of the IDP and that there is less sequence-dependent separation of *R_g_* between different IDPs with *λ*=1.10 than with the original force field (Fig. S9). Thus, a possible explanation for the worsened agreement with intrachain PREs for A2 LCD is that it is a relatively compact IDP (***Ryan et al., 2018***), and *λ*=1.10 may result in an overly expanded ensemble. This may also be the reason why our set of multidomain proteins seems to require a lower value of *λ* than IDPs do; two of the multidomain proteins in the set are variants of full-length A1 for which the isolated IDR requires only *λ*=1.04 for optimal agreement with SAXS. Thus, full-length A1 may be a similar outlier due to the properties of its IDR. Additionally, although the accuracy of IDP-IDP affinities is improved, the modified force field does not seem to accurately distinguish between IDPs that should and should not self-associate, as FUS LCD showed a similar level of self-association to IDPs that should not self-associate. Despite these limitations to our approach, the modified version of Martini 3 with protein-water interactions rescaled by *λ*=1.10 shows clear improvement over the original force field in capturing the global dimensions, interactions, and affinities of IDPs. We stress that this modification to the force field is only tested for proteins in solution, and may not be applicable to all classes of biomolecules in Martini. We also note that additional experimental studies that quantify weak, transient interactions between both highly soluble and more interaction-prone IDPs would be very useful to address these issues.

The issues discussed above illustrate that rescaling all protein-water interactions by a single factor *λ* is a somewhat ad-hoc approach to improve Martini 3, as the modified force field does not fully capture the sequence-dependent physico-chemical properties of proteins. Some of these issues could potentially be addressed by a more detailed reparameterization of Martini, for example by modifying the interactions of specific bead types or amino acids separately. To investigate which specific bead types or amino acids could require modified interaction potentials, we determined the correlation between the optimal value of *λ* for each protein and the bead type composition (Fig. S10) as well as other sequence metrics (Fig. S11). However, this approach did not uncover any clear strategy for reparameterizing specific bead types or amino acids. An alternative approach to favor more expanded conformations of IDPs could be the addition of IDR-specific backbone dihedrals, similar to the secondary structure-specific dihedrals already implemented in Martini. However, our results indicate that addition of dihedral potentials to capture for example transient secondary structure in IDPs would not solve the problems with overestimated compaction.

The functions of some IDPs and multidomain proteins depend on their ability to form biomolecular condensates (***Boeynaems et al., 2018***), often involving the formation of transient and multivalent protein-protein interactions and liquid-liquid phase separation (LLPS). Generally, the propensity of an IDP to undergo LLPS is correlated with its single-chain compactness (***Choi et al., 2020***). A modified version of Martini 2.2 with decreased protein-protein interactions has already been shown to improve the description of LLPS of an IDP (***Benayad et al., 2021***), and Martini 3 has also been used to study salt-dependent condensate formation (***Tsanai et al., 2021***). We expect that increased protein-water interactions, yielding improved accuracy of the global dimensions of IDPs and weakened IDP-IDP interactions, will be useful in future applications of Martini 3 to study the role of IDPs in biomolecular condensates as well astheirsingle-chain conformations and dynamics.

## Methods

### IDP simulations

We selected a set of twelve IDPs of varied sequence, with lengths between 24 and 334 amino acid residues and with SAXS data available: the N-terminal region of pertactin (PNt) (***Riback et al., 2017***), the NR interaction domain of N-CoR (CoRNID) (***Cordeiro et al., 2019***), two deletion mutants of Tau (K19 and K25) (***Mylonas et al., 2008***), the ‘plug domain from a TonB-dependent receptor (FhuA) (***Riback et al., 2017***), α-synuclein (aSyn) (***Ahmed et al., 2021***), the low-complexity domain of hnRNPA1 (A1) (***Martin et al., 2020***), theT-domain of colicin N (ColNT) (***Johnson et al., 2017***), Sic1 (***Gomes et al., 2020***), the activation domain of ACTR (ACTR) (***Kjaergaard et al., 2010***), Histatin-5 (Hst5) (***Jephthah et al., 2019***) and a tandem repeat of Histatin-5 (Hst52) (***Fagerberg et al., 2020***) (Table 1).

**Table 1.**
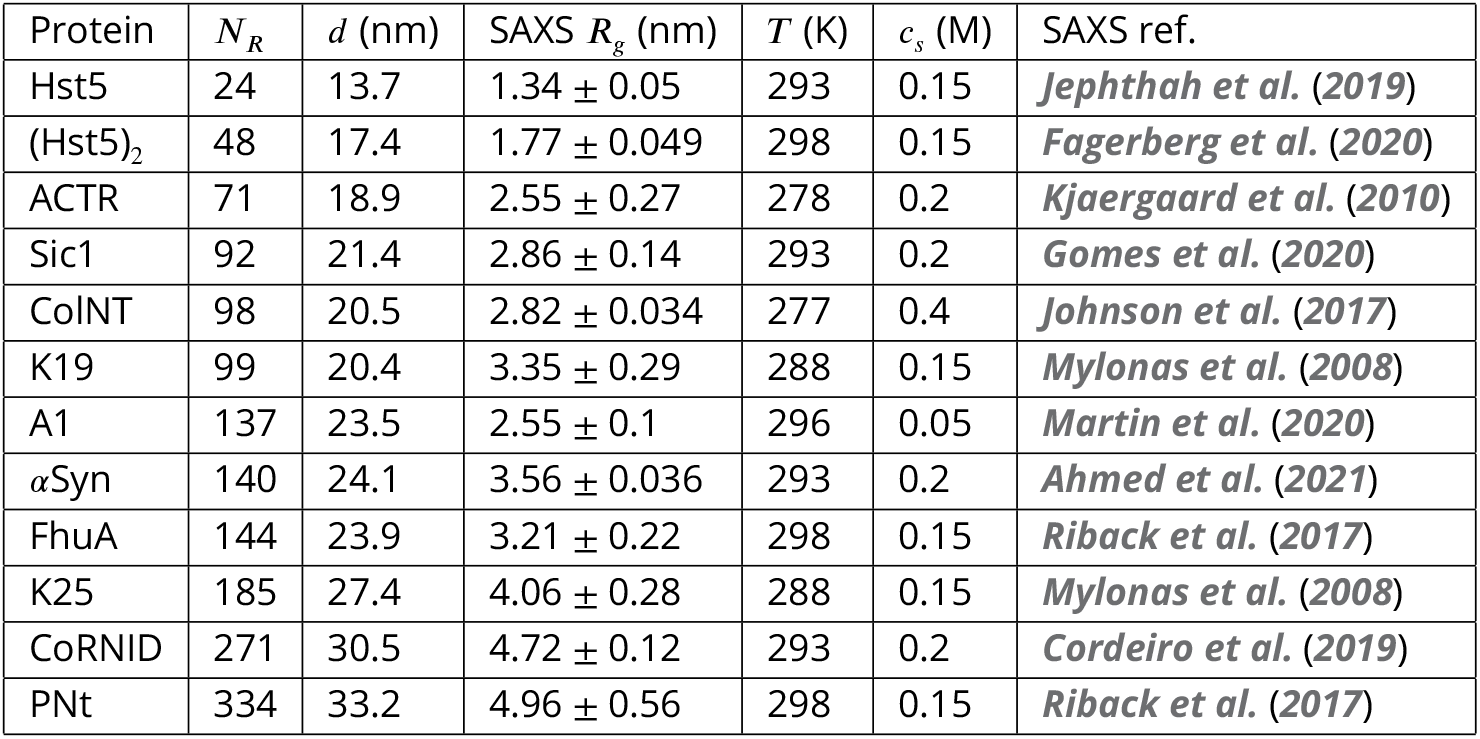
IDP simulations for SAXS and *R_g_* calculations: Number of amino acid residues (*N_R_*), box size (*d*), experimental *R_g_*, simulation temperature (*T*), and salt concentration in the simulation (*c_s_*).

We performed all MD simulations using Gromacs 2020.3 (***Abraham et al., 2015***) and the Martini 3.0 force field (***Souza et al., 2021***) or adapted force fields with rescaled protein-water interactions. Proteins were coarse-grained using the Martinize2 python script, placed in a dodecahedral box using Gromacs and solvated using the Insane python script (***Wassenaar et al., 2015***). Initial box sizes were chosen by using starting structures from simulations in ***Tesei et al. (2021b)*** corresponding to the 95th percentile of *R_g_*-distributions and using Gromacs *editconf* with the flag *-d 4.0*. Box sizes were later increased if necessary. NaCl concentration was set to match the conditions in SAXS experiments and to neutralize the system. No secondary structure or elastic network model was assigned with Martinize2 for IDPs and IDRs (see below for tests on ACTR). Energy minimization was performed using steepest descent for 10,000 steps with a 30 fs timestep. The Lennard-Jones potentials between all protein and water beads were rescaled by a factor *λ*. Seven values of *λ* were tested for each system: 1.00 (original force field), 1.04,1.06,1.08,1.10,1.12 and 1.14. The systems were equilibrated for 10 ns with a 2 fs timestep using the Velocity-Rescaling thermostat (***Bussi et al., 2007***) and Parinello-Rahman barostat (***Parrinello and Rahman, 1981***). Production simulations were run for between 27 μs and 40 μs with a 20 fs timestep using the Velocity-Rescaling thermostat and Parinello-Rahman barostat. The temperature was set to match conditions in SAXS experiments and the pressure was set to 1 bar. Non-bonded interactions were treated with the Verlet cut-off scheme. The cut-off for Van der Waals interactions was set to 1.1 nm. Coulomb interactions were treated using the reaction-field method with a 1.1 nm cut-off and dielectric constant of 15. Frames were saved every 1 ns. Periodic boundary conditions were treated with Gromacs *trjconv* with the flags *-pbc whole-center*. Simulation convergence was assessed using block-error analysis (***Flyvbjerg and Petersen, 1989***)of the *R_g_* using the BLOCKING code (https://github.com/fpesceKU/BLOCKING). Simulations were backmapped to all-atom using a modified (***Larsen et al., 2020***) version of the Backward algorithm (***Wassenaar et al., 2014**),* in which simulation runs are excluded and energy minimization is shortened to 200 steps.

We also ran MD simulations of five IDPs with paramagnetic relaxation enhancement (PRE) data available: the low-complexity domain of FUS (FUS) (***Monahan et al., 2017***), the low-complexity domain of hnRNPA2 (A2) (***Ryan et al., 2018***) aSyn (***Dedmon et al., 2005***), full-length tau (htau40) (***Mukrasch et al., 2009***) and osteopontin (OPN) (***Platzer et al., 2011***). For these proteins we set the NaCl concentration and temperature to match the conditions in PRE experiments. Simulations were otherwise set up and run using the same protocol as above.

### Multidomain protein simulations

We selected a set of three multidomain proteins with SAXS data available: full-length hnRNPA1 (FL-A1), full-length hnRNPA1 with an N-terminal His-SUMO tag (hSUMO-FL-A1) and TIA-1 (Table 2). SAXS data and initial structures of FL-A1 and hSUMO-FL-A1 were taken from ***Martin et al. (2021)***. These structures were built based on the structures of SUMO1 (PDB: 1A5R) (***Bayer et al., 1998***) and the RRM1 and RRM2 domains (PDB: 1HA1) (***Shamoo et al., 1997***). The initial structure of TIA-1 was taken from ***Larsen et al. (2020)*** and SAXS data was taken from ***Sonntag et al. (2017)***. The structure was built based on the structures of RRM1 (PDB 5O2V) (***Sonntag et al., 2017***), RRM2 (PDB: 5O3J) (***Sonntag et al., 2017***) and the RRM2-RRM3 complex (PDB: 2MJN) (***Wang et al., 2014***).

**Table 2.** Multidomain protein simulations for SAXS and *R_g_* calculations: Number of amino acid residues (*N_R_*), box size (*d*), experimental *R_g_*, simulation temperature (*T*), and salt concentration in the simulation (*c_s_*).

| Protein | $N_R$ | $d$ (nm) | SAXS $R_g$ (nm) | $T$ (K) | $c_s$ (M) | SAXS ref. |
| --- | --- | --- | --- | --- | --- | --- |
| TIA1 | 275 | 15.9 | $2.75 \pm 0.031$ | 300 | 0.1 | *Sonntag et al. (2017)* |
| FL-A1 | 314 | 25.6 | $3.12 \pm 0.022$ | 300 | 0.15 | *Martin et al. (2021)* |
| hSUMO-FL-A1 | 433 | 28.4 | $3.37 \pm 0.014$ | 300 | 0.1 | *Martin et al. (2021)* |

Simulations of multidomain proteins were set up and run using the same protocol as for the IDP simulations with a few exceptions: (i) Secondary structure was assigned with DSSP (***Kabsch and Sander, 1983***) in Martinize2. Disordered regions were assigned as coil. (ii) An elastic network model was applied with Martinize2 to keep folded domains intact. Interdomain elastic restraints and the elastic restraints in disordered regions and linker regions were removed. The elastic restraints consisted of a harmonic potential of 700 kJ mol^-1^ nm^-2^ between backbone beads within a 0.9 nm cut-off. (iii) Dihedrals between side chains and backbone beads were added based on the initial structures with the *-scfix* flag in Martinize2. These dihedrals were removed for disordered regions and linker regions. (iv) *λ*=1.02 was also tested. Simulations of FL-A1 and TIA1 were run for 40 μs. Simulations of hSUMO-FL-A1 were run for 15.9-19.4 μs.

### Simulations of dimerization of folded proteins

Initial structures of ubiquitin were taken from ***Vijay-Kumar et al. (1987)*** (PDB: 1UBQ). Initial structures of villin HP36 were taken from ***McKnight et al. (1997)*** (PDB: 1VII). Simulations of folded proteins were set up and run using the same protocol as for IDP simulations with a few exceptions: (i) Two copies of ubiquitin were placed in a cubic box with 14.92 nm sides, giving a protein concentration of 1 mM. Two copies villin HP36 were placed in a cubic box with 7.31 nm sides, giving a protein concentration of 8.5 mM. (ii) Secondary structure was assigned with DSSP (***Kabsch and Sander, 1983***) in Martinize2. (iii) An elastic network model was applied with Martinize2. The elastic restraints consisted of a harmonic potential of 700 kJ mol^-1^ nm^-2^ between backbone beads within a 0.9 nm cut-off. For ubiquitin, we removed elastic restraints from the C-terminus (residue 72-76) to allow for flexibility (***Lindorff-Larsen et al., 2005***). (iv) Dihedrals between side chains and backbone beads were added based on the initial structures with the *-scfix* flag in Martinize2. We ran simulations testing three different values of *λ*: 1.00,1.10, and 1.12. For each value of *λ*, we ran ten replicas of 40 μs each.

### Simulation of dimerization of IDPs

Simulations of two copies of FUS, aSyn, htau40, and p15_*paf*_ were set up and run using the same protocol as for IDP simulations. To match conditions in reference experiments ***(Monahan et al., 2017; Dedmon et al., 2005; Mukrasch et al., 2009; De Biasio et al., 2014***), the proteins were placed in cubic boxes with 40.5, 25.51,48.02, and 34.15 nm side lengths respectively, giving total protein concentrations of 50, 200, 30, and 83.4 μM. We ran simulations testing three different values of *λ*: 1.00,1.10, and 1.12, with ten replicas for each *λ*. Simulations of FUS, aSyn, htau40, and p15_*paf*_ were run for 11.5-21.0, 40, 7.7-12.7, and 16.2-35.8 μs respectively.

### IDP simulations to test the effect of long-range electrostatics, secondary structure, and more bulk solvent

To test whether inclusion of long-range electrostatics affects the compaction of IDPs, we ran simulations of Hst5 and aSyn using Particle Mesh Ewald (PME) electrostatics with a Fourier spacing of 0.16 nm, cubic interpolation, and a real-space cut-off of 1.1 nm. These simulations were otherwise set up and run using the same protocol as for other IDPs. Simulations of Hst5 were run for 20 μs and simulations of aSyn were run for 8-9.6 μs.

To test whether inclusion of secondary structure affects the compaction of IDPs, we ran simulations of ACTR with helix dihedrals applied to two transient helices at positions Glu28-Ser40 and Leu47-Leu54 (***Kjaergaard et al., 2010***). Assignment of helix dihedrals was performed with Martinize2. These simulations were otherwise set up and run using the same protocol as for other IDPs.

To test whether overestimated IDP compaction is the result of a lack of bulk solvent, we ran simulations of aSyn in a much larger box (*d*=34.1 nm). These simulations were otherwise set up and run using the same protocol as for other IDPs.

### Calculating the radius of gyration

We calculated the *R_g_* from the coarse-grained trajectories using Gromacs *gyrate* (***Abraham et al., 2015***). Figure S12 shows the distribution of *R_g_* for the 12 IDPs. We used block-error analysis (***Flyvbjerg and Petersen, 1989***) to estimate the error on the averages. Experimental *R_g_* and corresponding error bars were calculated from SAXS profiles by Guinier analysis using ATSASAUTORG with default settings (***Petoukhov et al., 2007***).

### SAXS calculations

After each trajectory had been backmapped to all-atom resolution, we extracted 15000 frames (evenly distributed in the time-series) to calculate SAXS profiles using Pepsi-SAXS (***Grudinin et al., 2017***). To avoid potential problems of overfitting the parameters for the contrast of the hydration layer (*δρ*) and the displaced solvent (*r*_0_) (if these are fitted individually to each structure) we used values that have previously been shown to provide good agreement with experiment for flexible proteins (***Pesce and Lindorff-Larsen, 2021***). Values for the intensity of the forward scattering (*I*(0)) and the constant background (*cst*) were fitted globally with least-squares regression weighted by the experimental errors using the Scikit-learn python library (***Pedregosa et al., 2011***).

To quantify the agreement between experimental SAXS profiles and those calculated from simulations, we calculated the reduced *χ*^2^:

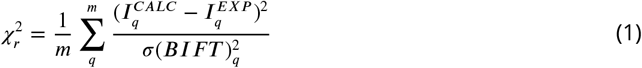

Here *m* is the number of data points, 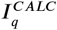 and 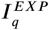 are the averaged calculated SAXS intensity and the experimental SAXS intensity at scattering angle *q*, and *σ(BIFT)_q_* is the error for the experimental intensity at scattering angle *q* corrected according to: 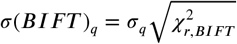, where *σ_q_* is the experimental error and 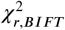 quantifies the agreement between the experimental SAXSdata and the model SAXS profile calculated from the pair distance distribution function obtained through the Bayesian Indirect Fourier Transform algorithm (BIFT) (***Hansen, 2000***). This approach has been shown to lead to improved error estimates for experimental SAXS profiles (***Larsen and Pedersen, 2021***) and, here, made it possible to compare more directly and average over the 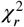 from the different systems. The BIFT algorithm optimizes the hyperparameter *D_max_* (maximum distance between scattering particles in the system) starting from an initial estimate. To set this initial estimate we, for each protein, used the largest value of *D_max_* observed over all simulations with different values of *λ*.

### PRE calculations

We used the DEER-PREdict software (***Tesei et al., 2021a***) to calculate PRE NMR data for three proteins (Table 3) from all-atom backmapped trajectories. DEER-PREdict implements a model-free formalism (***Iwahara et al., 2004***) combined with a rotamer library approach to describe the MTSL spin-label probe (***Polyhach et al., 2011***). We assumed an effective correlation time of the spin label, *τ_t’_* of 100 ps and scanned the molecular correlation time, *τ_c_*, from 1 to 20 ns in increments of 1 ns. Additionally, to calculate PRE intensity ratios, we assumed a transverse relaxation rate for the diamagnetic protein of 10 *s*^-1^ and approximated the total INEPT time of the HSQC measurement to 10 ms (***Battiste and Wagner, 2000***). We calculated intermolecular PRE rates from two-chain simulations treating one chain as spin-labeled and the other as ^15^N-labeled. We averaged the PRE rates obtained for the two combinations of spin-labeled and ^15^N-labeled chain. Agreement between calculated and experimental PREs was quantified by the reduced *χ*^2^ over all spin-label sites:

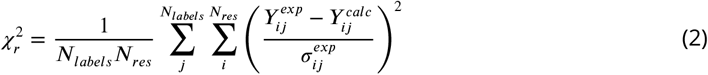

**Table 3.**
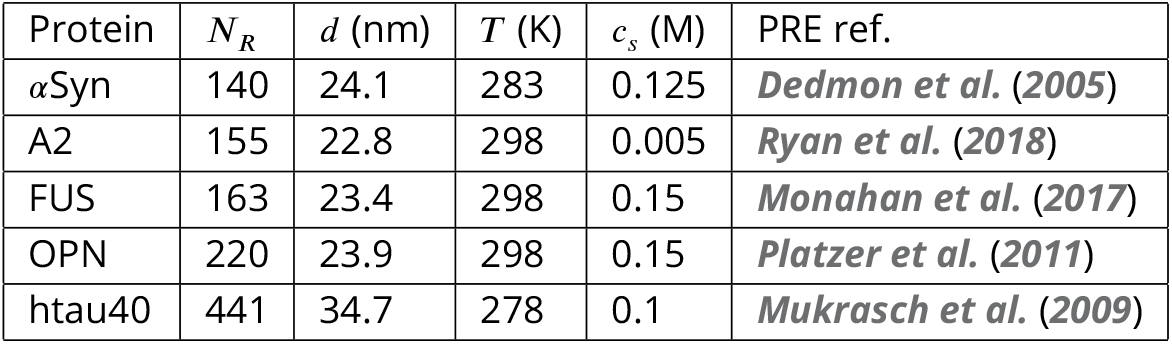
IDP simulations for single-chain PRE calculations: Number of amino acid residues (*N_R_*), box size (*d*), experimental *R_g_*, simulation temperature (*T*), and salt concentration in the simulation (*c_s_*).

| Protein | $N_R$ | $d$ (nm) | $T$ (K) | $c_s$ (M) | PRE ref. |
| --- | --- | --- | --- | --- | --- |
| αSyn | 140 | 24.1 | 283 | 0.125 | *Dedmon et al. (2005)* |
| A2 | 155 | 22.8 | 298 | 0.005 | *Ryan et al. (2018)* |
| FUS | 163 | 23.4 | 298 | 0.15 | *Monahan et al. (2017)* |
| OPN | 220 | 23.9 | 298 | 0.15 | *Platzer et al. (2011)* |
| htau40 | 441 | 34.7 | 278 | 0.1 | *Mukrasch et al. (2009)* |

Where *N_labels_* and *N_res_* are the number of spin-labels and residues, 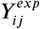 and 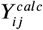 are the experimental and calculated PRE rates for label *j* and residue *i*, and 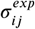 is the experimental error of the PRE rate for label *j* and residue *i*.

### Dimerization calculations

We analyzed the population of the bound and unbound states of ubiquitin, villin HP36, FUS, α-synuclein, htau40, and p15_*paf*_ homodimers in our simulations (Table 4). We calculated the minimum distance between any beads in the two proteins over the trajectory using Gromacs *mindist* (***Abraham et al., 2015***). The fraction bound was defined as the fraction of frames where the minimum distance was below 0.8 nm.

**Table 4.** Protein dimerization simulations: Number of amino acid residues (*N_R_*), box size (*d*), experimental *R_g_*, simulation temperature (*T*), and salt concentration in the simulation (*c_s_*).

| Protein | $N_R$ | $d$ (nm) | $T$ (K) | $c_s$ (M) | PRE or affinity ref. |
| --- | --- | --- | --- | --- | --- |
| αSyn | 140x2 | 25.51 | 283 | 0.125 | *Dedmon et al. (2005)* |
| FUS | 163x2 | 40.5 | 298 | 0.15 | *Monahan et al. (2017)* |
| p15PAF | 76x2 | 34.15 | 298 | 0.15 | *De Biasio et al. (2014)* |
| htau40 | 441x2 | 48.02 | 278 | 0.1 | *Mukrasch et al. (2009)* |
| ubq | 76x2 | 14.9 | 303 | 0.11 | *Liu et al. (2012)* |
| villin HP36 | 36x2 | 7.31 | 298 | 0.15 | *Brewer et al. (2005)* |

Forsimulationsof ubiquitin, the fraction bound was also calculated using the minimum distance only between beads in the binding site (residue 8,13,44,45,46,49,67,68,70,71, and 73) defined by NMR chemical shift perturbations (***Liu et al., 2012***). This greatly reduced population of the bound state, showing that Martini3 did not capture the specificity of the interaction. For ubiquitin and villin HP36 dimerization, we calculated what the fraction bound should be at the concentration in our simulations based on the *K_d_*-values of 4.9 mM and 1.5 mM respectively (***Liu et al., 2012; Brewer et al., 2005***). The fraction bound was calculated as:

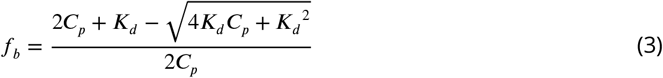

where *f_b_* is the fraction bound, *C_p_* is the concentration of one of the copies of the protein in the simulation box, and *K_d_* is the dissociation constant.

## Data availability

Scripts and data are available via http://github.com/KULL-Centre/papers/tree/main/2021/Martini-Thomasen-et-al

## Acknowledgments

We thank Simone Orioli, Thea K. Schulze and Yong Wang for useful discussions and suggestions. We acknowledge the use of computational resources from Computerome 2.0 and the corefacility for biocomputing at the Department of Biology. This research was supported by the Lundbeck Foundation BRAINSTRUC initiative (R155-2015-2666 to K.L.-L.).

