## Supporting Figures for "Improving Martini 3 for disordered and multidomain proteins"

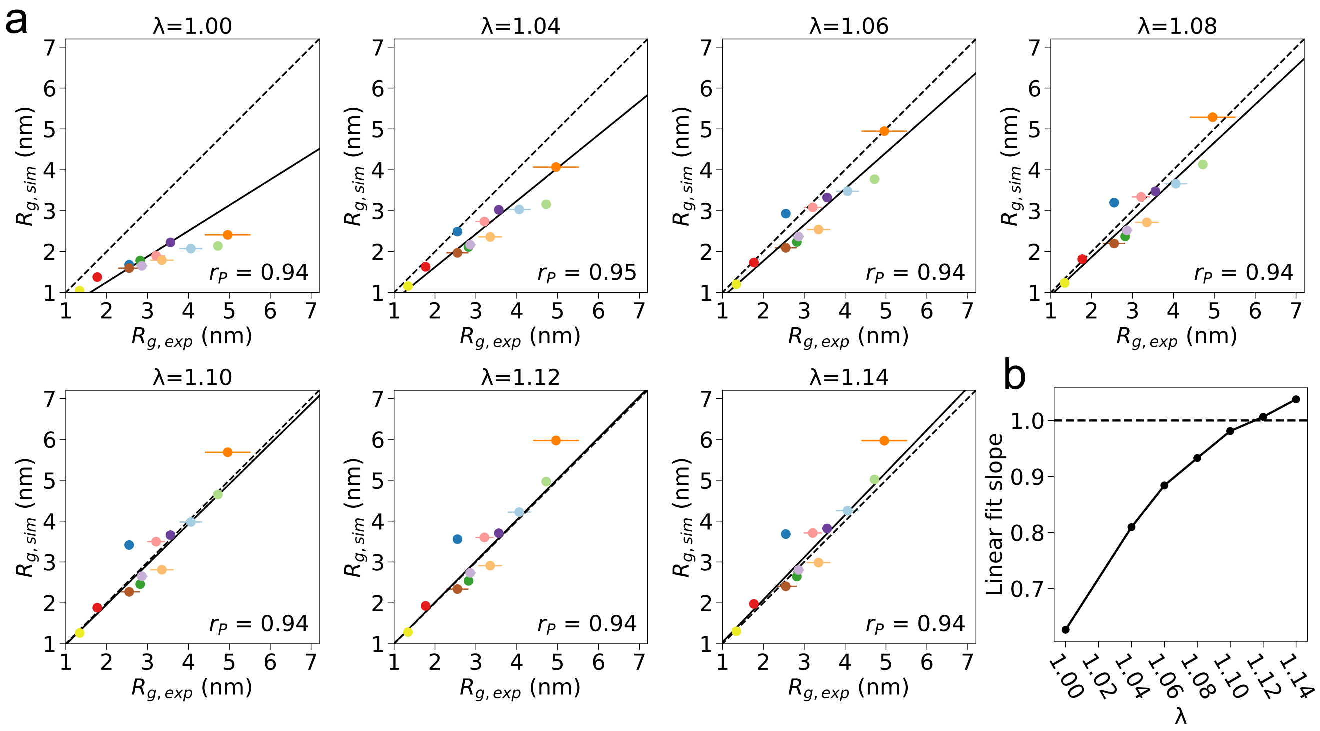


**Figure S1. Increased protein-water interactions improve the agreement with experimental $R_g$ for IDPs**
**a.** Average $R_g$ from MD simulations over a range of different protein-water interaction rescaling factors $\lambda$ plotted against experimental $R_g$ from Guinier analysis of SAXS data for a set of twelve IDPs. Experimental error bars from Guinier fit and simulation error bars determined by block error analysis (***Flyvbjerg and Petersen, 1989***) are shown. Linear fit with intercept 0 weighted by experimental errors is shown as a solid line. **b.** Slope of the linear fit shown in figure a with slope=1 marked by dashed line.

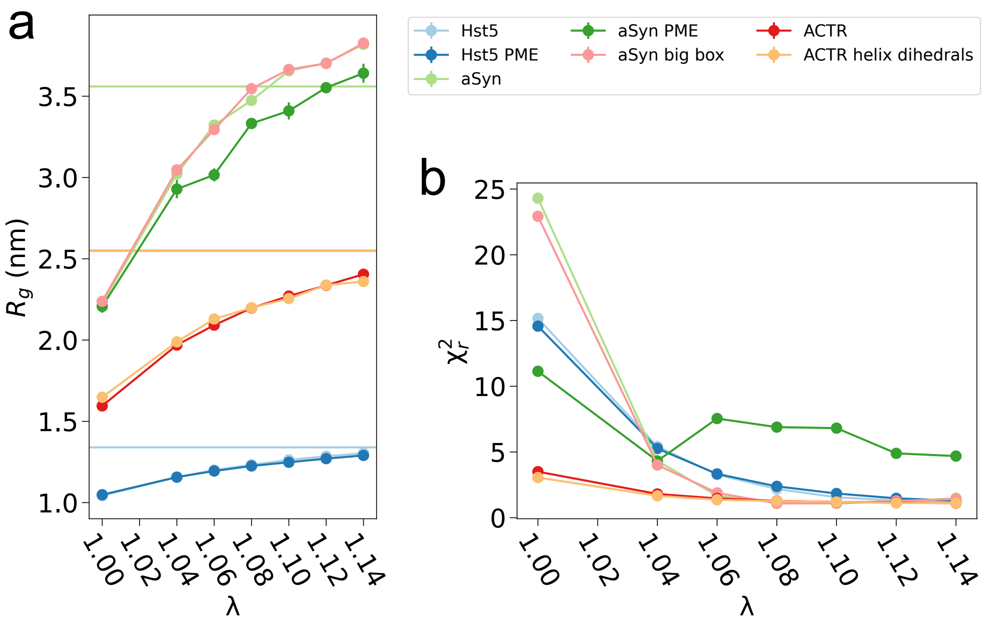


**Figure S2. a.** Average $R_g$ from MD simulations over a range of $\lambda$-values testing changes to the simulation setup. Experimental values from Guinier analysis of SAXS data are shown as horizontal lines. Simulations of Hst5 and aSyn were run with Particle Mesh Ewald (PME) electrostatics. Simulations of aSyn were run in a large box to increase the amount of bulk solvent. ACTR was run with helix dihedrals assigned at the positions of two transient helices (*Kjaergaard et al., 2010*). $R_g$-values from simulations run with the standard setup are also shown. **b.** Reduced $\chi_r^2$ between SAXS profiles calculated from MD simulations and experimental SAXS profiles for a range of $\lambda$-values for the same simulations as in panel a.

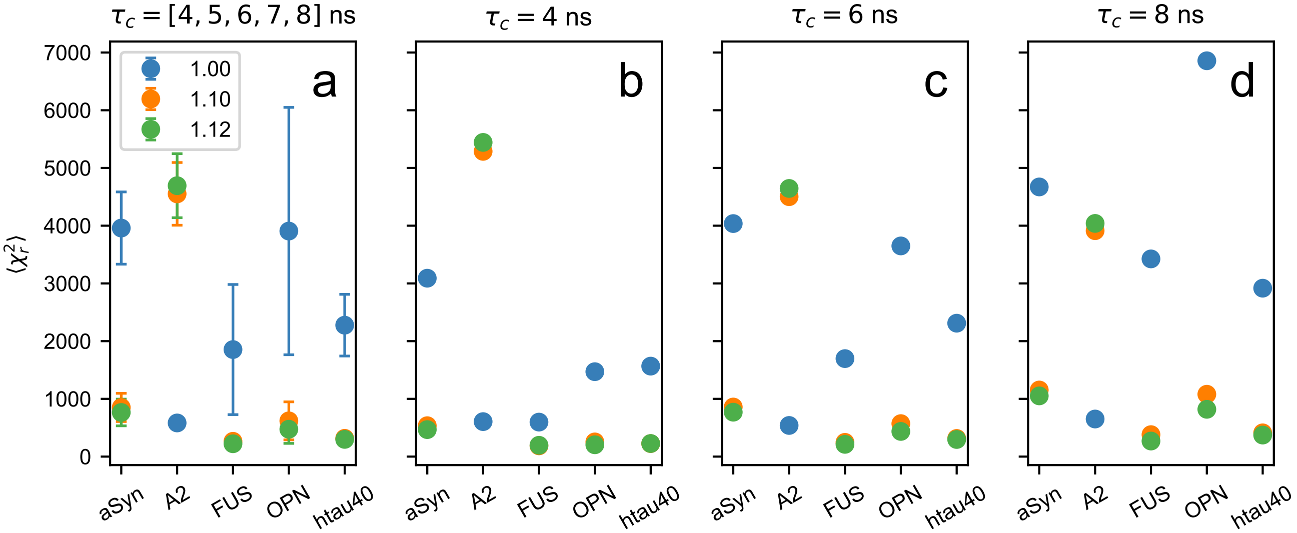


**Figure S3.** Effect of changing rotational correlation time $\tau_c$ in PRE calculations from simulations on the agreement with experimental PRE data ($\chi_r^2$). **A.** Average $\chi_r^2$ over PREs calculated with $\tau_c$ set to 4, 5, 6, 7, and 8 ns. Error bars show standard deviation. **B-D.** $\chi_r^2$ to experimental PREs with $\tau_c$=4, 6, and 8 ns.

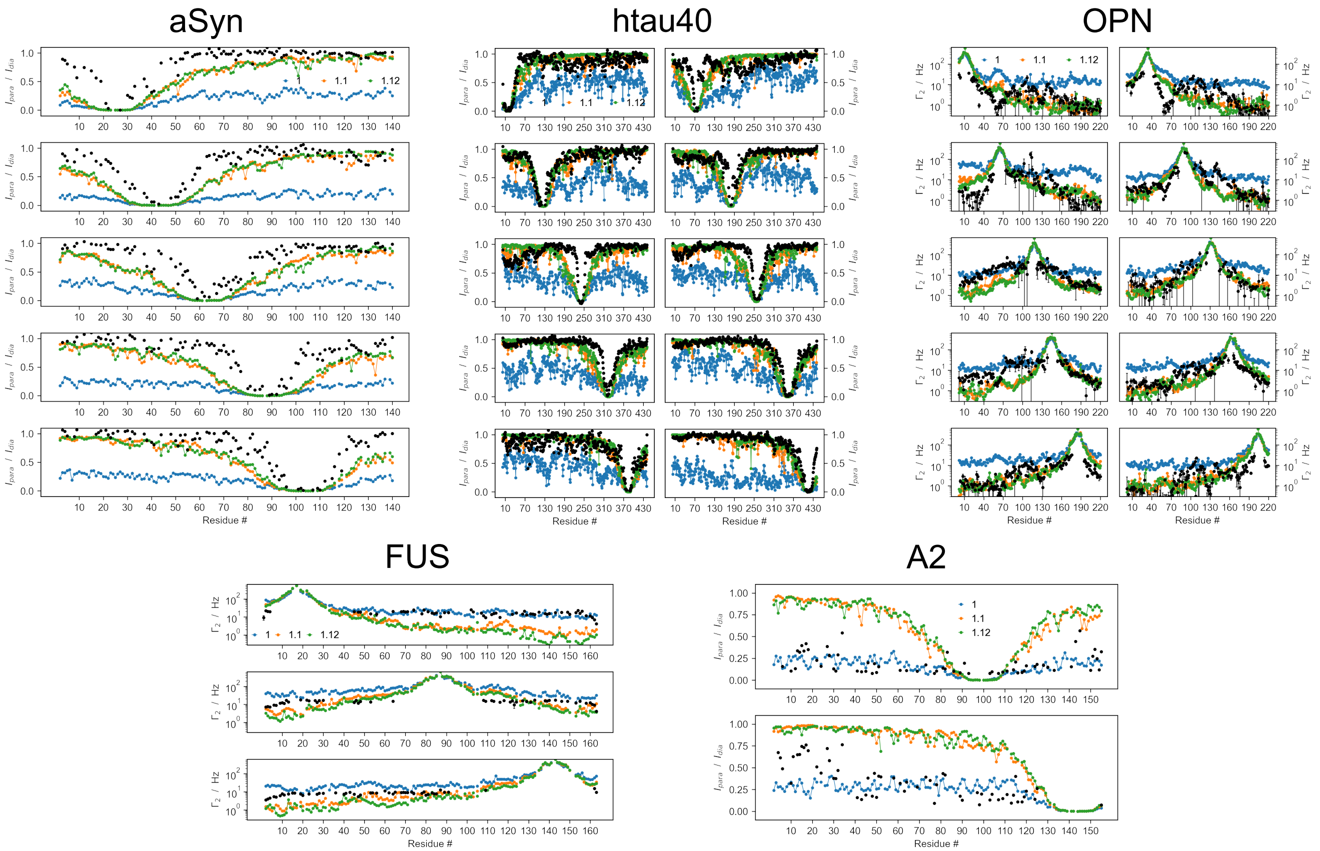


**Figure S4.** Paramagnetic relaxation enhancement (PRE) NMR data calculated from simulations of five IDPs with $\lambda$=1.00 (blue), $\lambda$=1.10 (orange), and $\lambda$=1.12 (green) with spin-labeling sites corresponding to experiments. $\tau_c$ was set to 4 ns. Experimental values are shown in black.

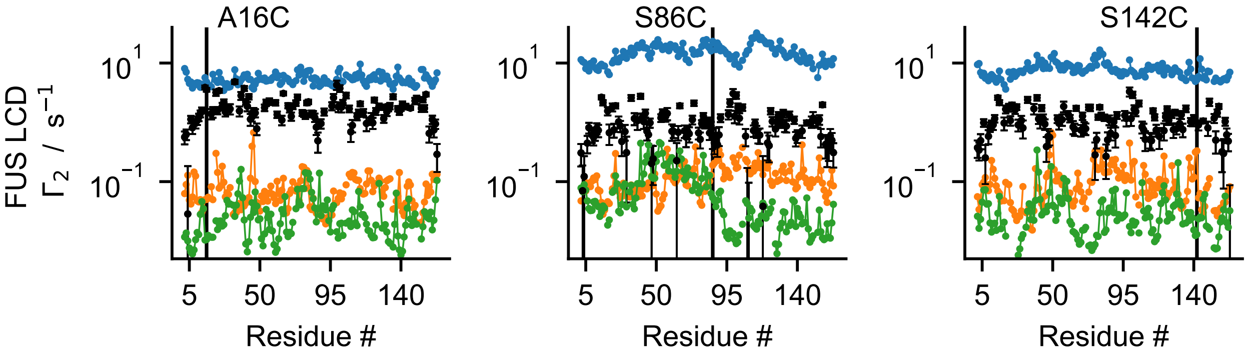


**Figure S5.** Intermolecular PREs calculated with $\tau_c$ set to 6 ns from simulations of two copies of FUS with $\lambda$=1.00 (blue), $\lambda$=1.10 (orange), and $\lambda$=1.12 (green). Experimental values are shown in black.

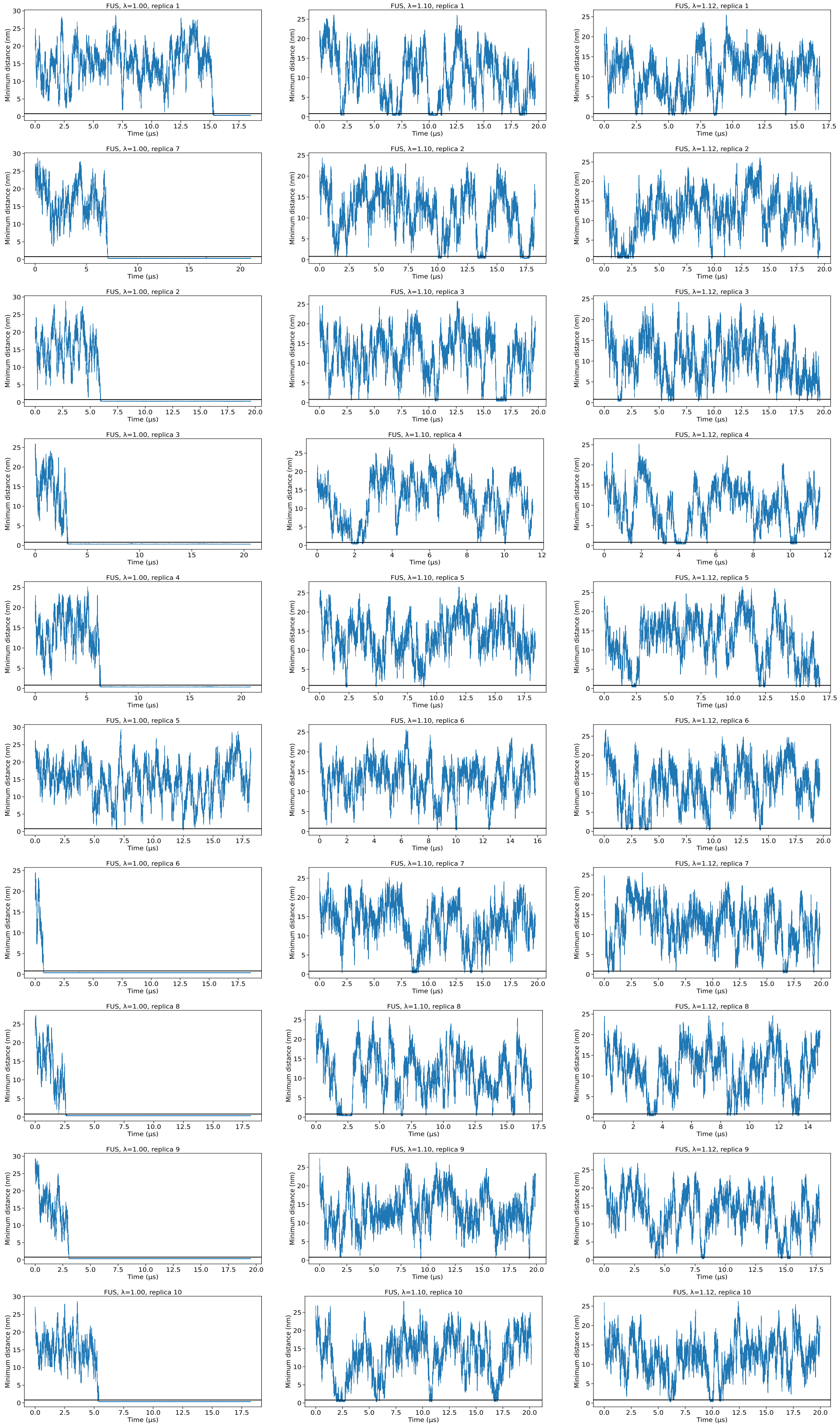


**Figure S6.** Time-series of the minimum distance between two copies of FUS LCD calculated from MD simulations with $\lambda$=1.00 (left), $\lambda$=1.10 (middle), and $\lambda$=1.12 (right).

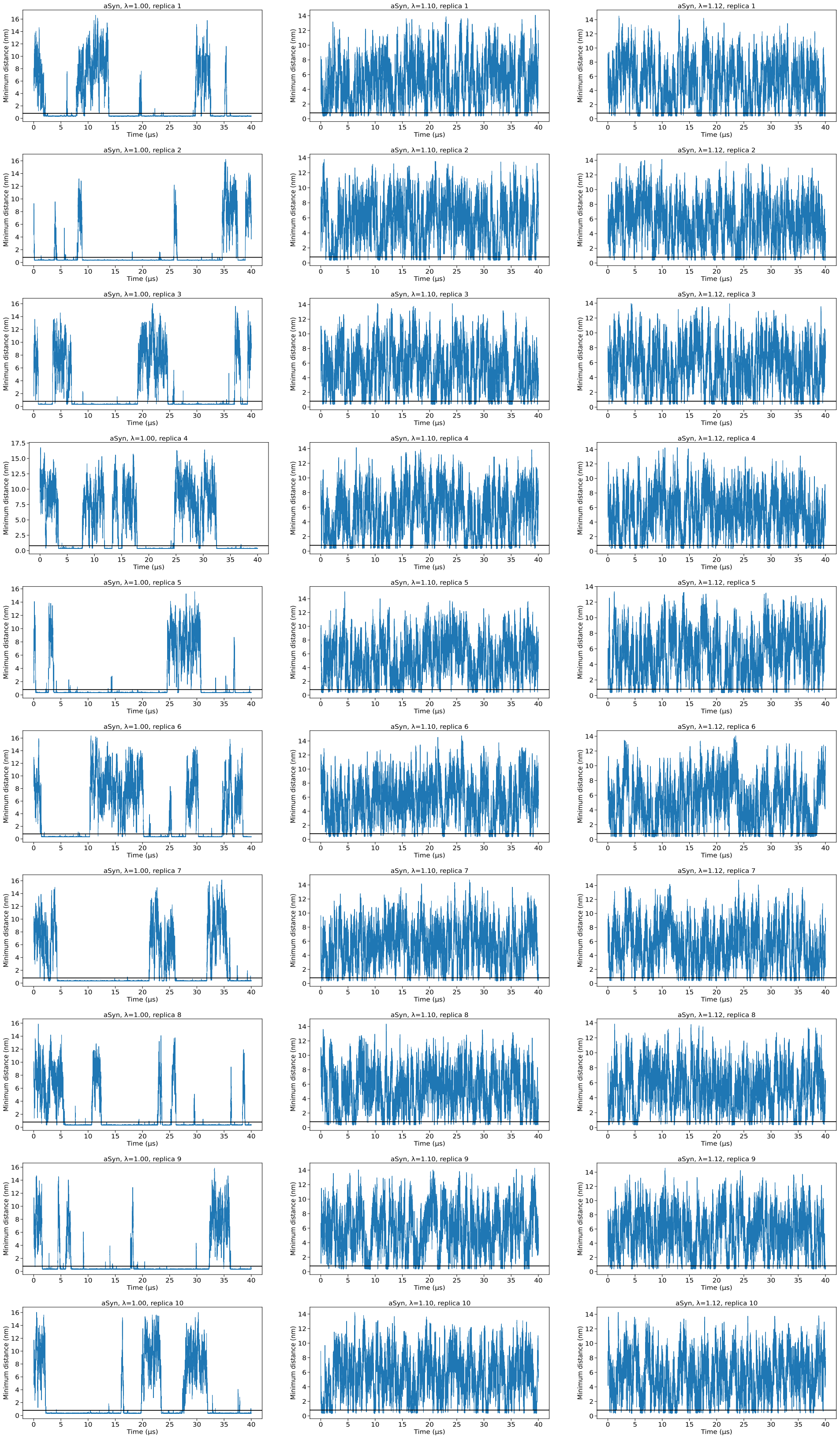


**Figure S7.** Time-series of the minimum distance between two copies of $\alpha$-synuclein calculated from MD simulations with $\lambda$=1.00 (left), $\lambda$=1.10 (middle), and $\lambda$=1.12 (right).

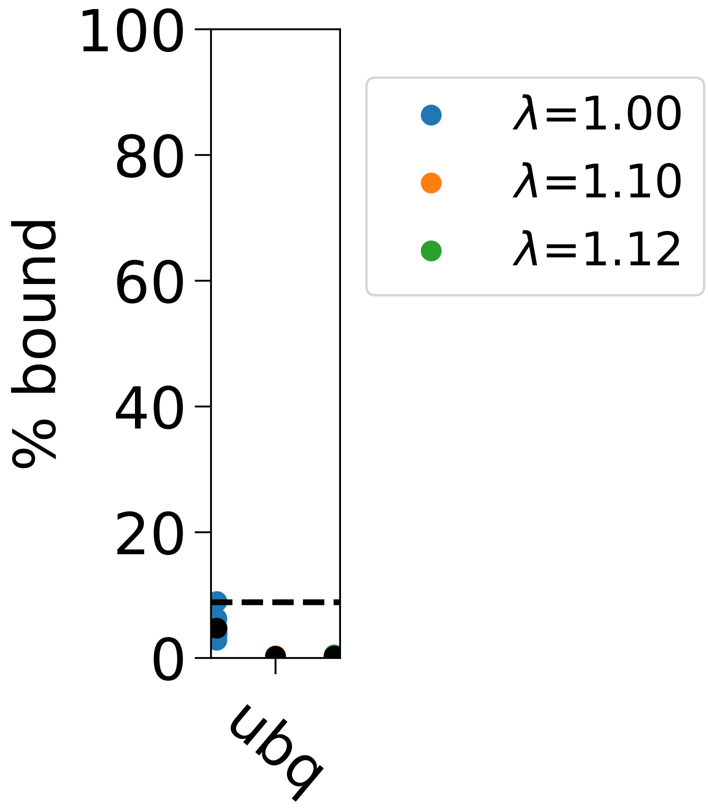


**Figure S8.** Fraction bound calculated from MD simulations of two copies of ubiquitin with different protein-water interaction rescaling factors $\lambda$. The bound state was defined using the minimum distance only between beads in the binding site determined by NMR chemical shift perturbations (***Liu et al., 2012***). The results from ten replica simulations are shown as colored points with the average value shown in black. The fraction bound in agreement with $K_d$=4.9mM for ubiquitin self-association is shown as a dashed line (***Liu et al., 2012***).

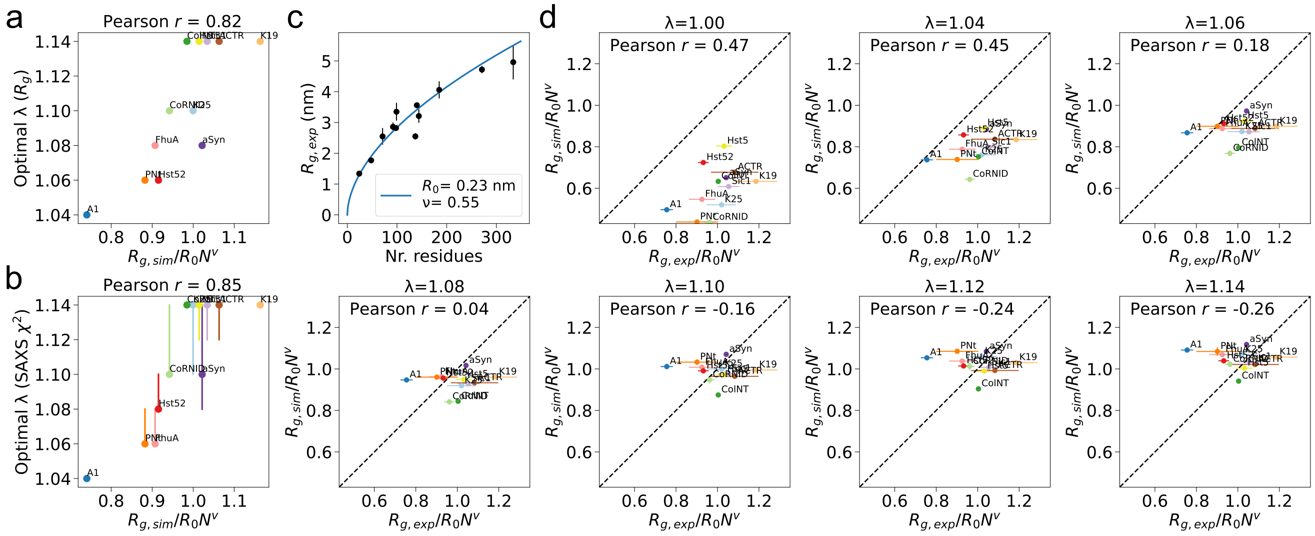


**Figure S9. Increased protein-water interactions reduce sequence-dependence of $R_g$**

**a.** Optimal $\lambda$ for each protein based on $R_g$ plotted against $R_g$ normalized by chain-length. **b.** Optimal $\lambda$ for each IDP based on $\chi_r^2$ to SAXS plotted against $R_g$ normalized by chain-length. Bars show the range of $\lambda$-values that give good agreement with SAXS data, selected as $\lambda$-values that give $\chi_r^2 < \chi_{min}^2 + (\chi_{max}^2 - \chi_{min}^2)0.02$, where $\chi_{min}^2$ and $\chi_{max}^2$ are the minimum and maximum $\chi_r^2$ given by any $\lambda$-value for the protein. **c.** Fit of $R_g$ power-law ($R_g = R_0 N^\nu$), where *N* is the number of residues, to experimental $R_g$ determined by Guinier analysis of SAXS for the set of twelve IDPs. The determined parameters were used to normalize $R_g$ to chain-length in figure b and c. **d.** Average $R_g$ from MD simulations with a range of protein-water interaction rescaling factors $\lambda$ normalized by chain-length plotted against experimental $R_g$ from Guinier analysis of SAXS data normalized by chain-length for a set of twelve IDPs.

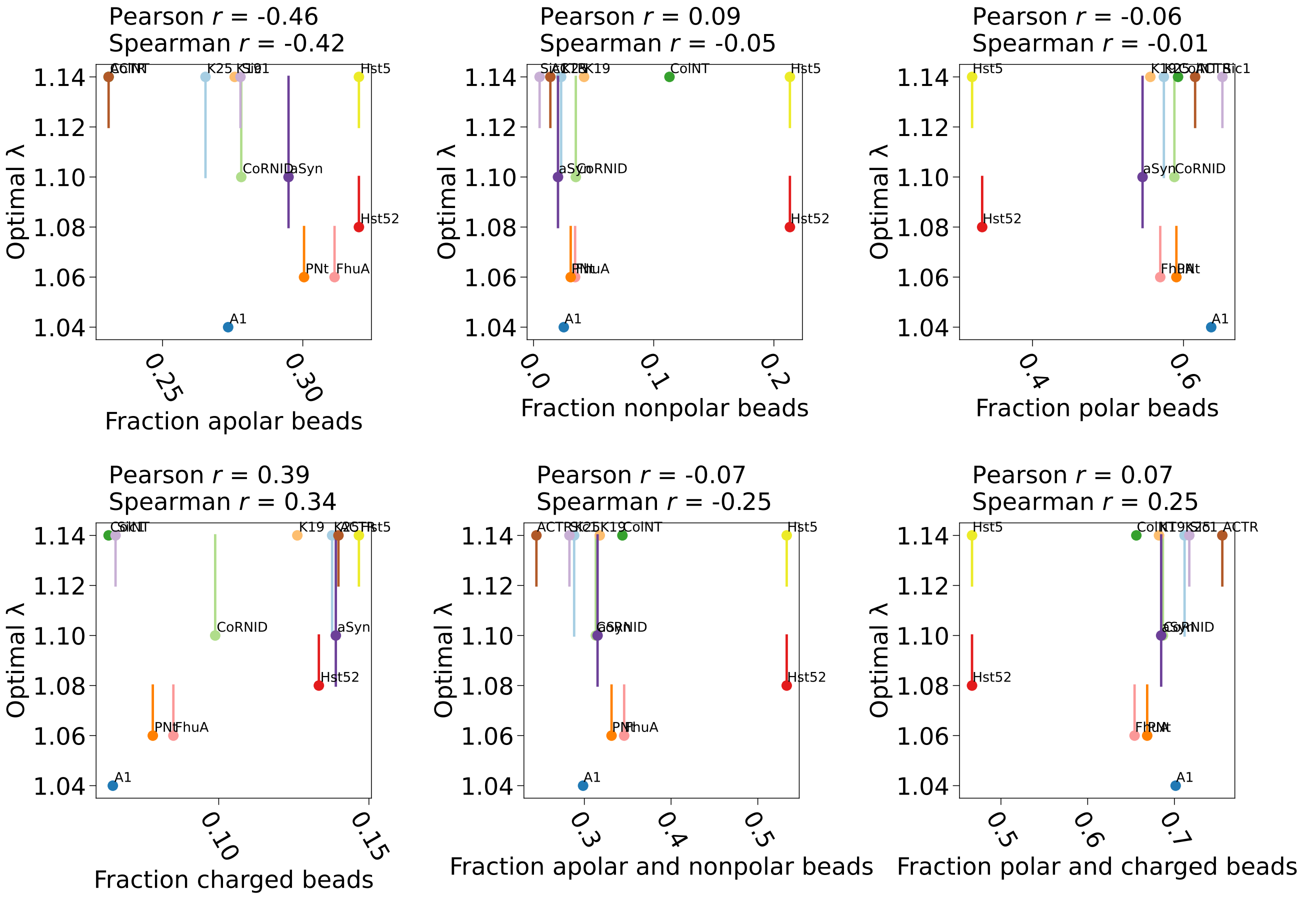


**Figure S10.** Optimal $\lambda$ determined for each IDP based on $\chi_r^2$ to SAXS plotted against the fraction of Martini bead types in the protein. Bars show the range of $\lambda$-values that give good agreement with SAXS data, selected as $\lambda$-values that give $\chi_r^2 < \chi_{min}^2 + (\chi_{max}^2 - \chi_{min}^2)0.02$, where $\chi_{min}^2$ and $\chi_{max}^2$ are the minimum and maximum $\chi_r^2$ given by any $\lambda$-value for the protein. Pearson and Spearman correlation coefficients are shown.

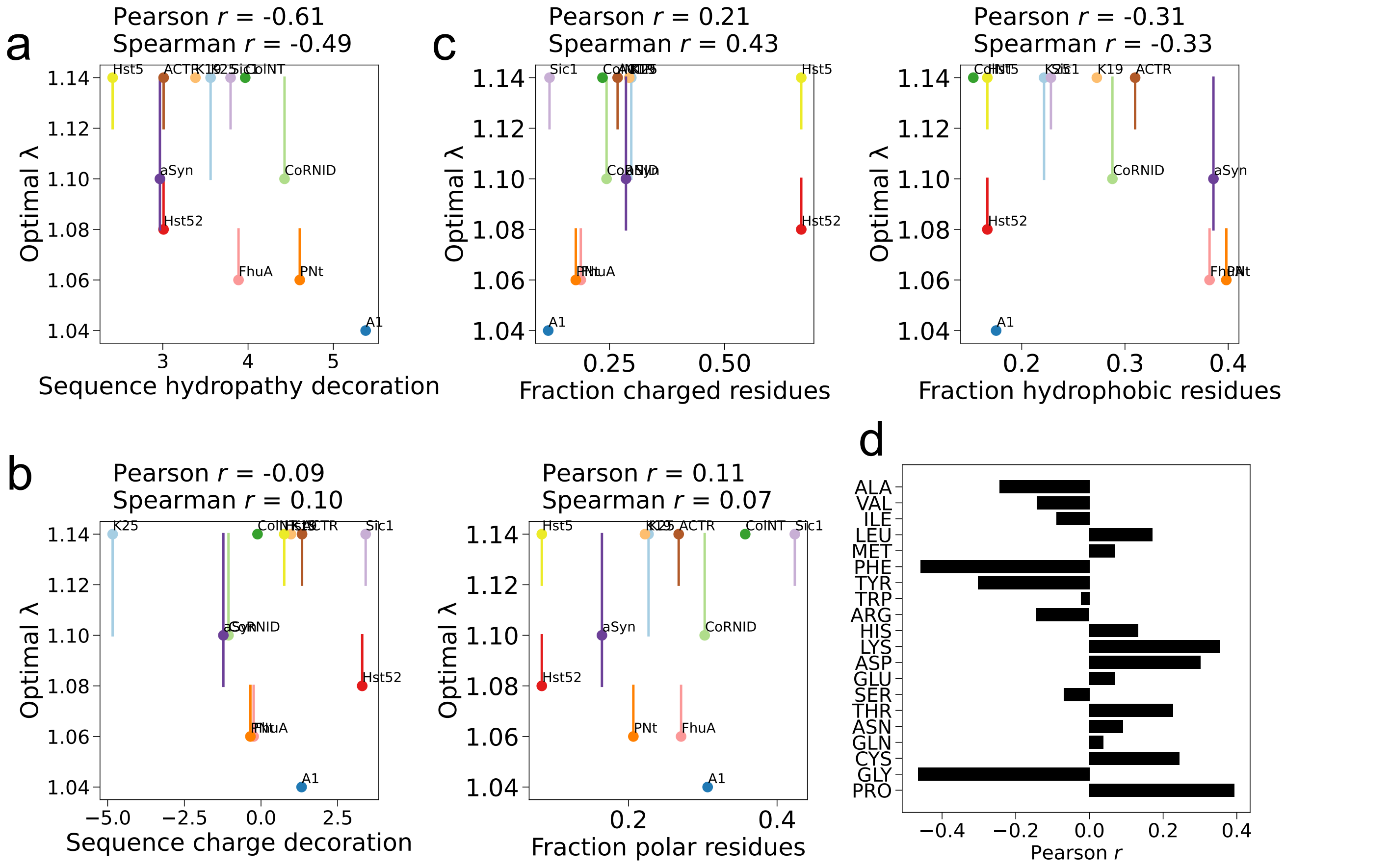


**Figure S11.** Optimal $\lambda$ determined for each IDP based on $\chi_r^2$ to SAXS plotted against different sequence metrics: **a.** Sequence hydropathy decoration (***Zheng et al., 2020***) using amino acid hydropathies from the M1 scale (***Tesei et al., 2021***). **b.** Sequence charge decoration (***Sawle and Ghosh, 2015***) **c.** Amino acid characteristics. Bars show the range of $\lambda$-values that give good agreement with SAXS data, selected as $\lambda$-values that give $\chi_r^2 < \chi_{min}^2 + (\chi_{max}^2 - \chi_{min}^2)0.02$, where $\chi_{min}^2$ and $\chi_{max}^2$ are the minimum and maximum $\chi_r^2$ given by any $\lambda$-value for the protein. Pearson and Spearman correlation coefficients are shown for each plot. **d.** Pearson correlation coefficients for correlation between optimal $\lambda$ determined for each IDP based on $\chi_r^2$ to SAXS and fraction of amino acid type in the sequence.

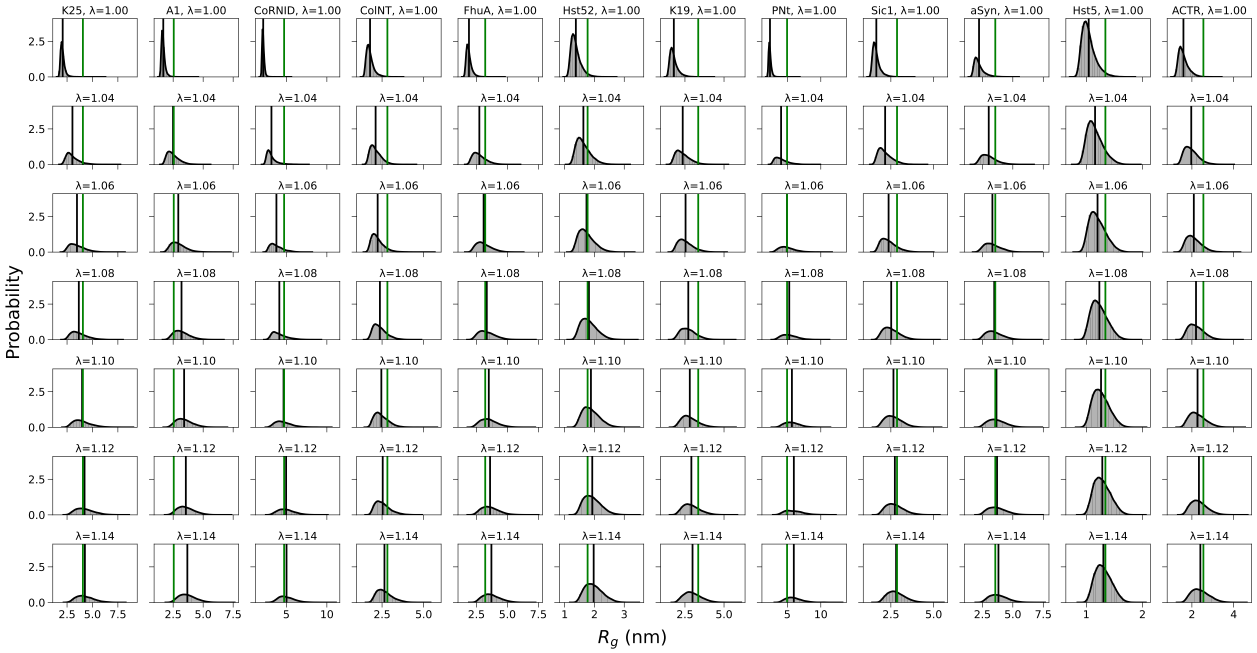


**Figure S12.** $R_g$-distributions from MD simulations over a range of protein-water interaction rescaling factors $\lambda$ for a set of twelve IDPs. Average is shown in black. Experimental values from Guinier analysis of SAXS data are shown in green.

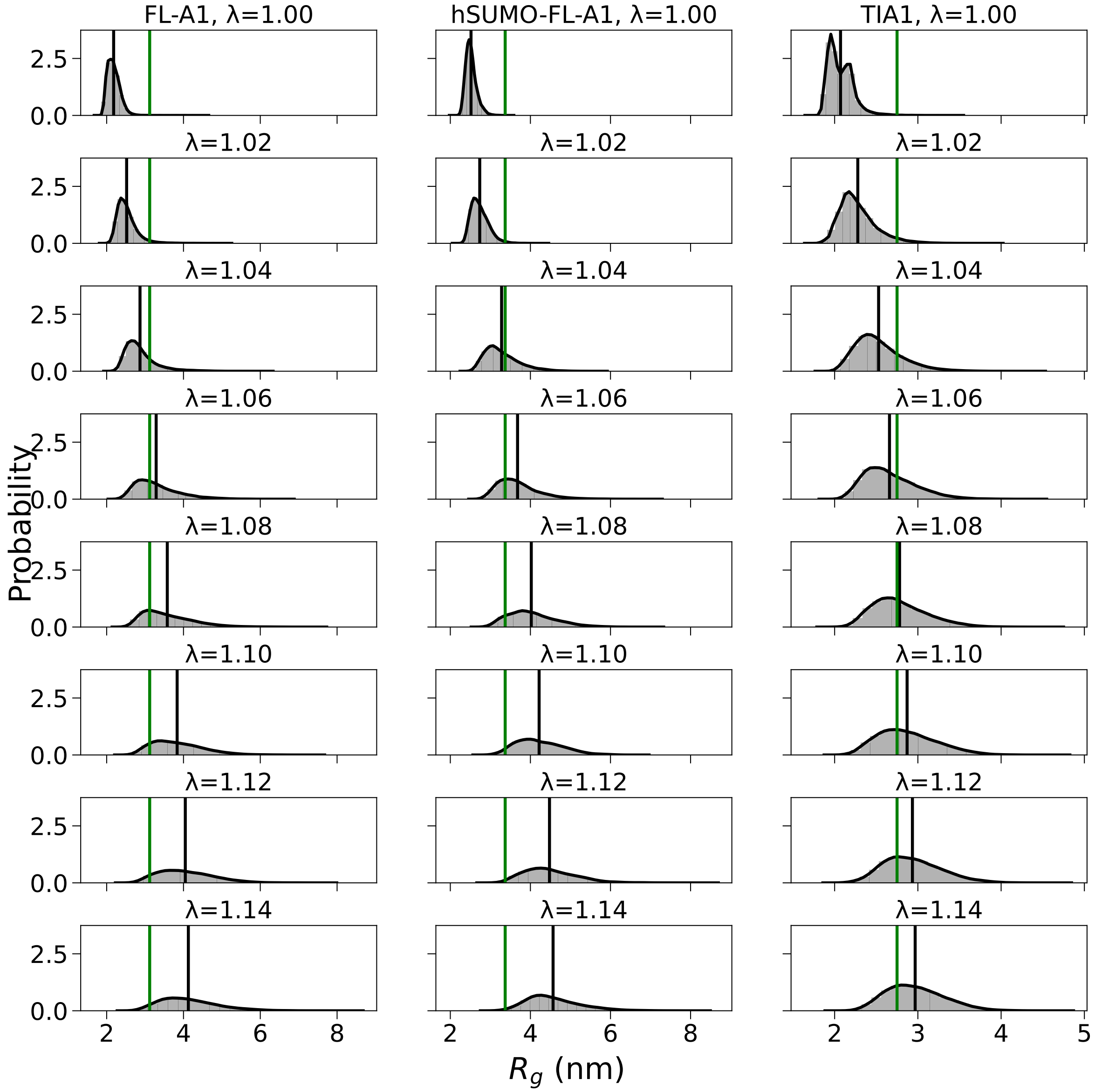


**Figure S13.** $R_g$-distributions from MD simulations over a range of protein-water interaction rescaling factors $\lambda$ for three multidomain proteins. Average is shown in black. Experimental values from Guinier analysis of SAXS data are shown in green.

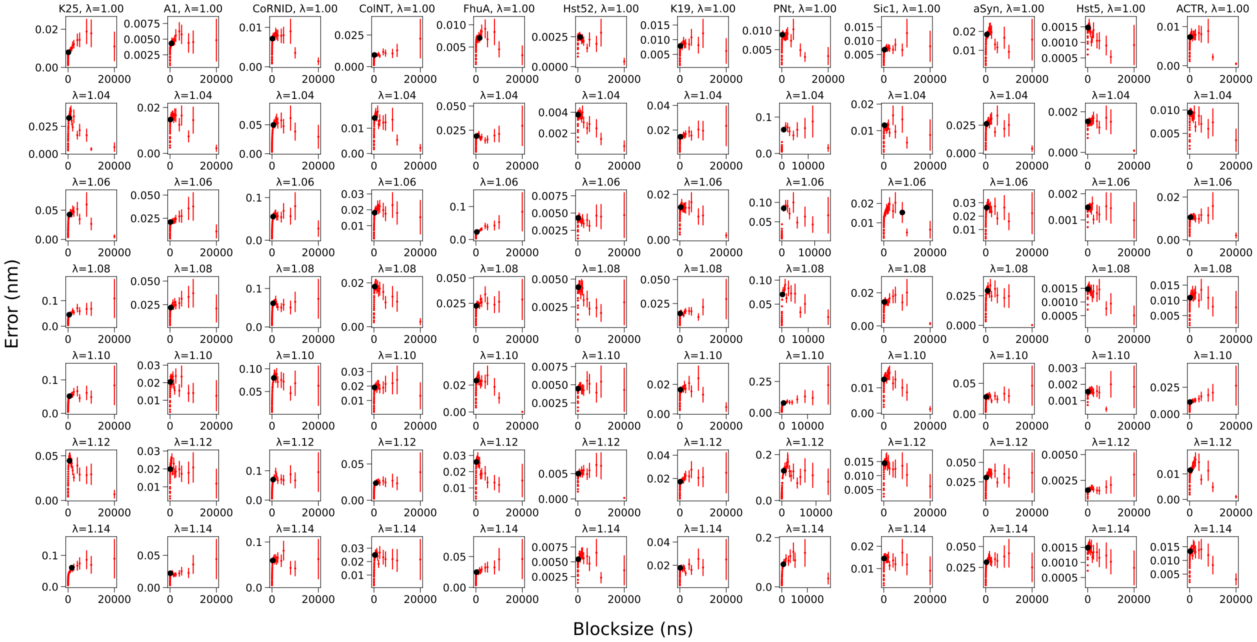

**Figure S14.** Block error analysis of the $R_g$ time-series from MD simulations of the twelve IDPs (***Flyvbjerg and Petersen, 1989***). The selected block size and error is shown in black.

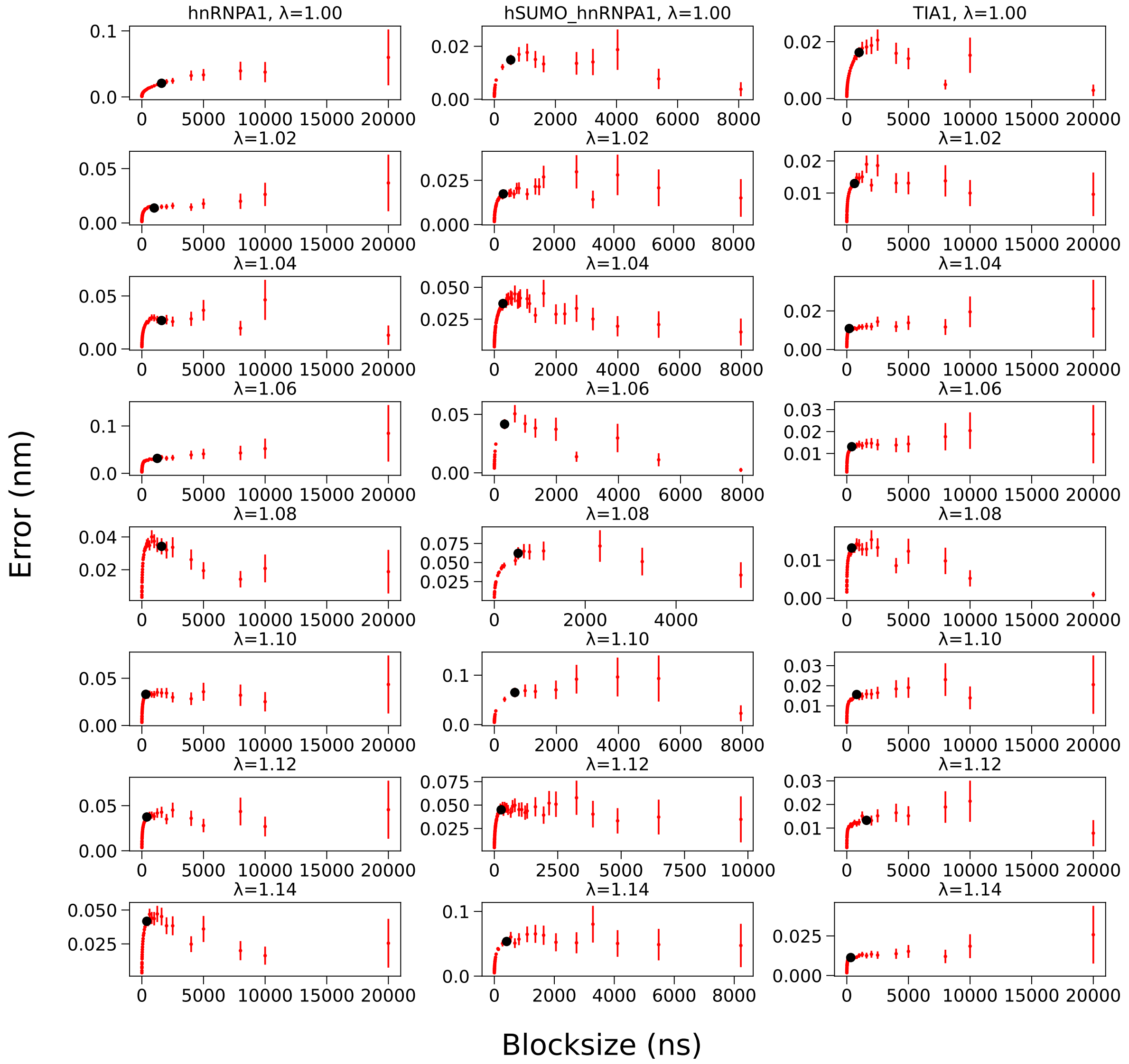


**Figure S15.** Block error analysis of the $R_g$ time-series from MD simulations of the three multidomain proteins (***Flyvbjerg and Petersen, 1989***). The selected block size and error is shown in black.
